# Spaced training forms complementary long-term memories of opposite valence in *Drosophila*

**DOI:** 10.1101/785618

**Authors:** Pedro F. Jacob, Scott Waddell

**Affiliations:** Centre for Neural Circuits & Behaviour, University of Oxford, Oxford, OX1 3TA, United Kingdom

## Abstract

Forming long-term memory (LTM) in many cases requires repetitive experience spread over time. In *Drosophila*, aversive olfactory LTM is optimal following spaced training, multiple trials of differential odor conditioning with rest intervals. Studies often compare memory after spaced to that after massed training, same number of trials without interval. Here we show flies acquire additional information after spaced training, forming an aversive memory for the shock-paired odor and a ‘safety-memory’ for the explicitly unpaired odor. Safety-memory requires repetition, order and spacing of the training trials and relies on specific subsets of rewarding dopaminergic neurons. Co-existence of the aversive and safety memories can be measured as depression of odor-specific responses at different combinations of junctions in the mushroom body output network. Combining two particular outputs appears to signal relative safety. Learning a complementary safety memory thereby augments LTM performance after spaced training by making the odor preference more certain.

## Introduction

The ability to learn and remember is fundamental for animals to adapt to a constantly changing environment, allowing them to anticipate meaningful events and to respond appropriately to specific stimuli. Across the animal kingdom, forming robust long-term memory (LTM) often requires multiple training trials with intervening rest periods, or intertrial intervals (ITI) (Ebbinghaus, 1913; Carew et al., 1972; Tully et al., 1994; Kogan et al., 1997; Hermitte et al., 1999; Menzel et al., 2001). Prevailing models suggest that optimal interval timing coincides with the dynamics of cellular signalling processes that are essential for LTM (Zhang et al., 2012; Liu et al., 2013; Smolen et al., 2016)

In *Drosophila*, LTM formation is maximal following 5 to 10 spaced training trials with a 15 min ITI, where an individual training trial pairs one of two odors with an electric shock reinforcement. In contrast, repeating the same number of trials, but without an ITI, referred to as massed training, forms a distinct consolidated form of memory referred to as ARM, Anaesthesia Resistant Memory (Tully et al., 1994). A large number of studies have uncovered molecular mechanisms that differentiate between ARM and LTM formed after either massed or spaced training. For example, flies mutant for the *radish* (*rad*) gene have been reported to specifically lack aversive ARM, whereas pharmacological and genetic blockers of new transcription and protein synthesis only disrupt LTM (Tully et al., 1994; Yin et al., 1994; Dubnau et al., 2003; Chen et al., 2012; Miyashita et al., 2012). The 15 min ITI required for optimal spaced training coincides with the peak of training-induced activity of the extracellular signal-regulated kinase (ERK, aka MAPK) (Pagani et al., 2009; Miyashita et al., 2018). In parallel with mechanisms of plasticity in other species, activated ERK phosphorylates and activates gene expression driven by the cAMP-response element binding (CREB) transcription factor (Bartsch et al., 1998; Impey et al., 1998; Thomas and Huganir, 2004; Miyashita et al., 2018). In *Drosophila* spaced training activated CREB induces expression of the c-Fos transcription factor, encoded by the *kayak* gene. c-Fos in turn is required to activate CREB and a positive CREB-cFos feedback loop prolongs the increased CREB activity that is essential to sustain LTM (Miyashita et al., 2018). In the mouse elevated CREB activity appears to provide an eligibility trace - it increases the likelihood that neurons become part of a memory engram (Han et al., 2007; Zhou et al., 2009; Park et al., 2016). Consistent with this model, spaced training produces more c-Fos positive Kenyon Cells (KCs) in the fly mushroom body (MB), and blocking output from all c-Fos labelled neurons impairs expression of LTM (Miyashita et al., 2018).

Research in *Drosophila* has also provided a cellular and neural circuit context for memory formation and retrieval. Subsets of anatomically restricted dopaminergic neurons (DANs) provide reinforcement signals that modulate connections between MB Kenyon Cells (KCs) and MB output neurons (MBONs), whose dendrites occupy the same MB compartment (Claridge-Chang et al., 2009; Aso et al., 2010; Burke et al., 2012; Liu et al., 2012; Lin et al., 2014). DAN activity coincident with odor exposure depresses synaptic connections between sparse populations of odor-activated KCs and MBONs (Séjourné et al., 2011; Hige et al., 2015; Owald et al., 2015; Perisse et al., 2016) via dopamine receptor directed cAMP-dependent plasticity (Yu et al., 2006; Kim et al., 2007; Tomchik and Davis, 2009; Qin et al., 2012; Zhang and Roman, 2013; Boto et al., 2014; Hige et al., 2015). Aversive learning reduces odor-drive to approach generating MBONs, which primarily occupy the vertical MB lobe. In contrast, appetitive learning reduces responses of avoidance directing MBONs, mainly on the tips of the horizontal MB lobes. Memory formation therefore establishes different configurations of the MBON network and the trained odors subsequently drive the skewed output (Séjourné et al., 2011; Hige et al., 2015; Owald et al., 2015; Perisse et al., 2016). Aversive LTM expression after spaced training strongly relies on αβ KCs and on downstream vertical lobe MBONs (MB-V2, aka MBON-α2sc, MBON-α’3m and MBON-α’3p, and MB-V3 aka MBON-α3; Tanaka et al., 2008; Aso et al., 2014a) that pool outputs from the vertical α collaterals of αβ KCs (Pascual and Préat, 2001; Isabel et al., 2004; Yu et al., 2006; Séjourné et al., 2011; Huang et al., 2012; Bouzaiane et al., 2015). However, γ and α’β’ KCs have also been implicated in LTM, either directly or by virtue of a requirement for downstream MBONs, such as MBON-γ3, MBON-γ3β’1 and MBON-M4β’2mp (Akalal et al., 2010; Wu et al., 2017). Interestingly, the network requirements for aversive LTM expression are evidently different to that for expression of 24 h ARM (Bouzaiane et al., 2015; Wu et al., 2017). However, although ARM and LTM differ at the molecular and network levels, it is not clear if flies acquire comparable information, or memory content, after spaced and massed training.

Prior studies have demonstrated that flies can simultaneously, or sequentially, form parallel avoidance and approach memories, that compete to guide memory-directed behaviour (Das et al., 2014; Aso and Rubin, 2016; Felsenberg et al., 2017, 2018). In addition, initial memory performance is additive if one odor is paired with shock and the other odor with sugar, during conditioning (Tempel et al., 1983). Theoretically, the differential nature of massed and spaced training could give flies the opportunity to learn that the shock-paired odor (CS+) is to be avoided, whereas the non-reinforced odor (CS-) is safe. Here we show that aversive spaced training permits flies to learn more information than massed training. Flies form an odor-specific avoidance memory for the previously shock-paired odor and learn that the non-shocked odor is safe and can be approached. Flies only learn that the second odor is safe if it is presented after the shock-paired odor in each trial. Safety memory formation requires the activity of two classes of rewarding DANs. Parallel aversive and safety memories can be recorded as depression of odor-specific responses in a distributed collection of unique MBONs. In addition, plasticity of MBON-γ3β’1, MBON-γ3 connections is required for flies to assess relative safety. LTM performance after spaced training therefore arises from addition of complementary odor-specific avoidance and approach memories.

## Results

### Spaced training forms two memories of opposite valence

Learning in different ways produces memories of distinct duration. In *Drosophila* investigators frequently compare the differences in memory formed following a number of training trials with or without inter-trial intervals. Although 5-10 differential odor-shock training trials forms memory that can be measured 24 h later, the mechanistic molecular and network underpinnings are clearly different if training is spaced or massed (Tully et al., 1994). We therefore investigated whether distinguishable neural correlates of spaced training might arise from the flies learning different information than when taught with massed trials.

We conditioned flies to form aversive olfactory LTM using a spaced training protocol of 6 differential training cycles separated by 15 min intervals (**Figure 1A**). In agreement with prior studies this regimen induced a persistent LTM measurable at 24 h after training. Flies selectively avoided the previously shock paired odor (CS+) when given the choice between that odor and the previously non-reinforced odor (CS-). Interestingly, if the flies were instead tested for their preference between the CS+ and a novel odor, flies exhibited a significantly reduced memory performance, in comparison with the CS+ *vs*. CS- condition (one-way ANOVA: p<0.05; n=8-13). Moreover, if the flies were tested for their preference between the CS- and a novel odor, they showed significant approach to the CS- (one sample t-test: p<0.05, significantly different from zero; n=10). These data are consistent with spaced training forming an avoidance memory for the CS+ and an approach memory for the CS- odor. These memories together contribute to the performance observed after spaced training.

**Figure 1.**
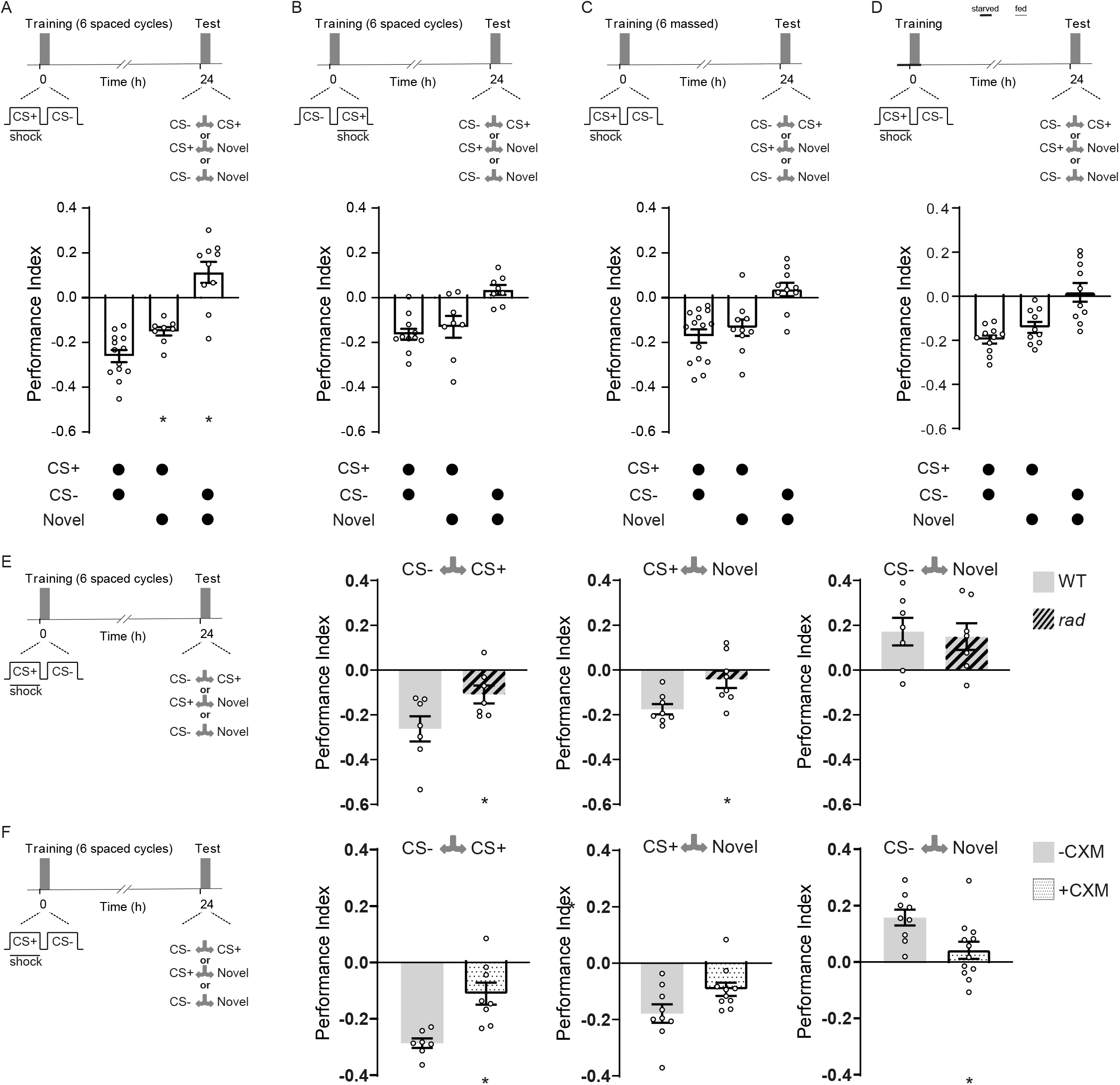
Spaced training induces LTM comprised of complementary CS+ and CS- components. (**A**) Spaced training (6 cycles of CS+/CS- training with 15 min ITI) generates LTM measured at 24 h. An aversive memory was measured at 24 h, when CS+ was tested against novel odor. An appetitive memory was measured at 24 h, when CS- was tested against novel odor. (**B** and **C**) If the order of CS+ and CS- were reversed during training (**B**) or there were no intervals between training cycles, massed training (**C**), the LTM (CS+ *vs*. CS-) was not different from the aversive memory of the CS+ (CS+ *vs*. Novel) and no approach was observed for the CS- (CS- *vs*. Novel). (**D**) A fasting LTM protocol that lacks repetition did not generate an appetitive memory to the CS- or a significantly different CS+ memory. (**E**) *Rad* mutant flies (hashed bars) displayed normal CS- approach memory but lacked evidence of CS+ memory in comparison to WT flies (grey bars). (**F**) CXM feeding impairs LTM performance after spaced training. Wild-type flies fed 5% glucose solution (grey bars) or glucose laced with 35 mM CXM (stippled bars) for 12-16 h overnight before spaced training. Asterisks denote significant differences. Data represented as mean ± standard error of mean (SEM). Individual data points displayed as dots.

In each cycle of the standard spaced protocol, the presentation of the CS+ precedes the CS-, which in principle could allow the flies to recognise that the CS- is not reinforced and perhaps ‘safe’. To challenge this mechanism, we reversed the order of CS+ and CS- presentation so that in each trial the CS- instead came before the CS+ (**Figure 1B**). Following the reversed spaced protocol flies displayed a CS+ *vs*. novel odor memory that was indistinguishable from the CS+ *vs*. CS- performance (one-way ANOVA: p>0.05; n=8-11). In addition, no CS- *vs*. novel odor performance was evident (one sample t-test: p>0.05, not significantly different from zero; n=8). The order of CS+ before CS- is therefore essential for flies to form a CS- approach memory after spaced training.

Given the reported difference in the cellular requirements between spaced and massed training, we tested whether 6 cycles of CS+/CS-massed training formed CS+ and CS- memories (**Figure 1C**). Mass trained flies did not show evidence for the formation of a CS- memory when tested CS- *vs*. novel odor (one sample t-test: p>0.05, not significantly different from zero; n=10). Consistent with a lack of CS- memory, their CS+ *vs*. novel odor performance was indistinguishable from CS+ vs. CS- memory (one-way ANOVA: p>0.05; n=10-15). Intervals between training trials are therefore also critical for the formation of CS- approach memory.

Hunger seems to change the rules for the formation of aversive LTM, so that a single training trial is more effective (Hirano et al., 2013). We therefore also tested the nature of memory formed following a ‘fasting LTM’ protocol (**Figure 1D**). Flies were starved for 12-16 h and then subjected to one round of CS+/CS- aversive training. After training flies were fed and memory performance was subsequently tested 24 h later. Although flies displayed significant 24 h memory performance we did not observe evidence for the formation of a CS- approach memory, when flies were tested for their preference between CS- and novel odor (one sample t-test: p>0.05, not significantly different from zero; n=10). Moreover CS+ *vs*. novel odor performance was indistinguishable from that flies tested with CS+ *vs*. CS- (oneway ANOVA: p>0.05; n=10-11). These data indicate that multiple trials are essential to form complementary CS+ avoidance and CS- approach memories.

### CS+ but not CS- memory is impaired in *radish* mutant flies

Seminal work concluded that spaced training forms protein synthesis dependent LTM, whereas massed training forms protein synthesis independent but *radish*-dependent consolidated ARM (Tully et al., 1994; Isabel et al., 2004). Since radish (*rad*) mutation (Folkers et al., 1993, 2006) is reported to specifically impair aversive ARM, we tested *rad* mutant flies for CS+ and CS- memory formation (**Figure 1E**). After spaced training, *rad* mutants showed impaired LTM performance (unpaired t-test: p<0.05; n=7). Surprisingly, *rad* mutant flies displayed normal CS- approach memory (unpaired t-test: p>0.05; n=7) but lacked evidence of CS+ memory (unpaired t-test: p<0.05; n=8). We also tested performance of flies fed with cycloheximide before training (**Figure 1F**). As previously reported, feeding the flies with cycloheximide (CXM) 12-16 h before spaced training (Tully et al., 1994; Yin et al., 1994) impaired LTM performance when flies were tested for their preference between the CS+ and CS- odors (unpaired t-test: p<0.001; n=7-8). Whereas CXM fed flies lacked CS- memory (unpaired t-test: p<0.05; n=9-12), aversive CS+ memory performance was not significantly altered (Mann Whitney test: p=0.054; n=9-10).

### Formation of CS- approach memory requires rewarding dopaminergic neurons

Several studies have established that DANs in the protocerebral anterior medial (PAM) cluster can provide reward-specific teaching signals during learning (Burke et al., 2012; Liu et al., 2012; Lin et al., 2014) (**Figure 2A**). We therefore tested whether their output was required to form the CS- appetitive memory after aversive spaced training (**Figure 2**). We expressed the dominant negative temperature sensitive UAS-*Shibire*^ts1^ (*Shi*^ts1^) encoded dynamin (Kitamoto et al., 2001) in rewarding DANs using R58E02-GAL4. Output from R58E02 DANs was specifically blocked during spaced training by raising the temperature of flies from 23°C to 32°C. Flies were then returned to 23°C and later tested for 24 h memory performance (**Figure 2A**). This manipulation impaired LTM performance when flies were tested for preference between the CS+ and CS- odors (one-way ANOVA: p<0.0001; n=14-15). Flies tested for preference between CS+ and novel odor revealed that aversive CS+ memory was relatively unaffected (**Figure 2B**, one-way ANOVA: p>0.05; n=6). However, no performance was evident when flies were tested for preference between CS- and novel odor (**Figure 2C**, one-way ANOVA: p<0.0001; n=7-8). Blocking rewarding DANs therefore specifically impaired formation of the CS- approach memory.

**Figure 2.**
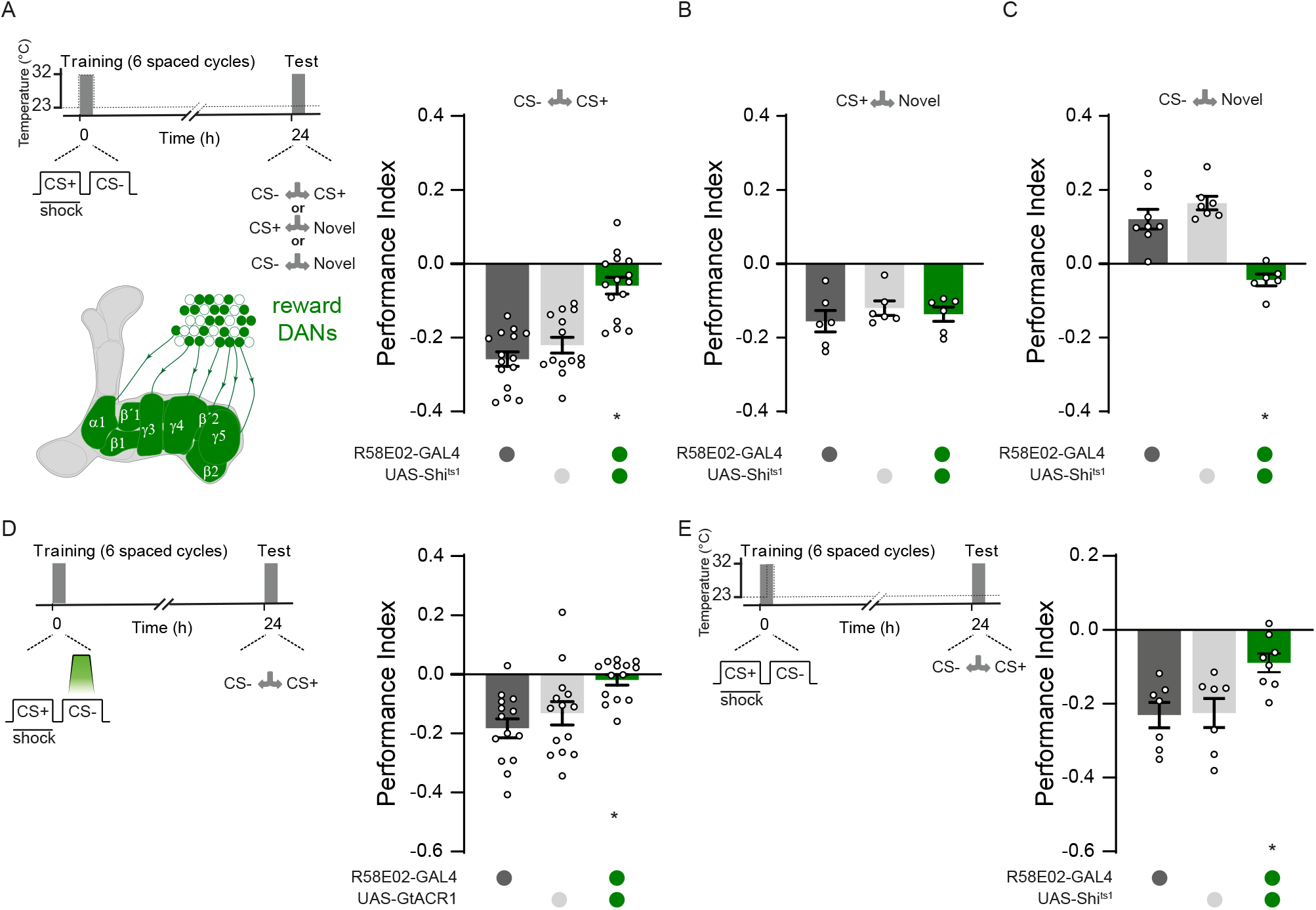
CS- approach memory requires PAM DANs during CS- presentation. (**A**) Left: protocol with temperature shifting (dashed line) and schematic of PAM DANs. Right: blocking PAM DANs with R58E02-GAL4; UAS-*Shi*^*t*s1^ during spaced training impairs 24 h memory performance. (**B**) Aversive memory to the CS+ is not affected. (**C**) Blocking PAM DANs specifically impairs, generating a slight aversion to the CS-. (**D**) Left: protocol with green light exposure during CS- presentation. Right: Blocking PAM DANs with R58E02-GAL4; UAS-GtACR1 during CS- impairs 24 h memory. (**E**) Left: protocol with temperature shifting (dashed line). Right: Blocking PAM DANs with R58E02-GAL4; UAS-*Shi*^ts1^, during the last three trials of spaced training, significantly reduces 24 h memory performance. Asterisks denote significant difference. Data represented as mean ± SEM. Individual data points displayed as dots. See also Figure S1.

We next used the light-gated GtACR1 anion channel (Mohamed et al., 2017) to restrict the timing of DAN inactivation specifically to the window of CS- presentation during each training cycle. This manipulation caused a similar impairment to 24 h CS+ *vs*. CS- performance (one-way ANOVA: p<0.05; n=14) as blocking DANs throughout the entire spaced training program (**Figure 2D**). These data are consistent with PAM DANs being required to reinforce the CS- approach memory during aversive spaced training.

We also tested the importance of trial repetition by using UAS-*Shi*^ts1^ to block R58E02 neurons during the last three trials of a six trial spaced training session (**Figure 2E**). Immediately following the third trial flies were transferred from 23°C to 32°C where training resumed. On termination of the sixth trial flies were then returned to 23°C and tested for 24 h CS+ vs. CS- LTM performance. This manipulation only significantly impaired memory performance of R58E02-GAL4; UAS-*Shi*^ts1^ flies demonstrating a requirement for PAM DAN output during the last three training trials (one-way ANOVA: p<0.05; n=7-8).

### CS+ avoidance and CS- approach memories co-exist in the MBON network

Prior studies have reported plasticity of odor-specific responses in the MB-V2 (MBON-α2sc) and MB-V3 (MBON-α3) MBONs as a correlate of aversive LTM after spaced training. The α2sc-MBONs exhibit a reduced response to the CS+ after aversive training and their output is required for the expression of aversive LTM (Séjourné et al., 2011). Plasticity and the role of α3-MBONs is more contentious. Pai et al. (2013) reported α3-MBONs to be required to express aversive LTM and an increased response to the CS+ response after spaced and not massed aversive training. However, Plaçais et al. (2013) reported an increased CS+ response only after appetitive training and for α3-MBON output to be dispensable for retrieval of aversive LTM. In contrast, appetitive memories, such as those reinforced by sugar, induce a relative depression of CS+ odor-evoked responses that can be measured in processes of the horizontal lobe M4β’ (MBON-β’2mp) and M6 (MBON-γ5β’2a) MBONs (Owald et al., 2015). Lastly, aversive memory can be extinguished by forming a parallel appetitive memory that manifests as a reduced CS+ odor-evoked response in MBON-γ5β’2a (Felsenberg et al., 2018).

We therefore used *in vivo* calcium imaging to test for odor-evoked physiological correlates of the CS+ aversive and CS- approach memories in these MBONs after spaced training. Ca^2+^ imaging was performed 24 h after training, so flies were trained in the T-maze and captured and mounted briefly before imaging.

We first tested whether we could reproduce previously reported physiological correlates of aversive LTM in the vertical lobe α2sc and α3-MBONs. Consistent with prior work we observed a significantly reduced CS+ odor-evoked response in the dendritic field of α2sc-MBONs after spaced training (**Figure 3A**). However, contrary to both prior reports, we also observed very robust depression of CS+ odor-evoked responses when recording in the dendrites of α3-MBONs (**Figure 3B**). Decreased CS+ responses in both types of vertical lobe MBONs after aversive spaced training is consistent with memory-directed CS+ odor avoidance arising from reduced α2sc and α3-MBONs mediated odor-approach.

**Figure 3.**
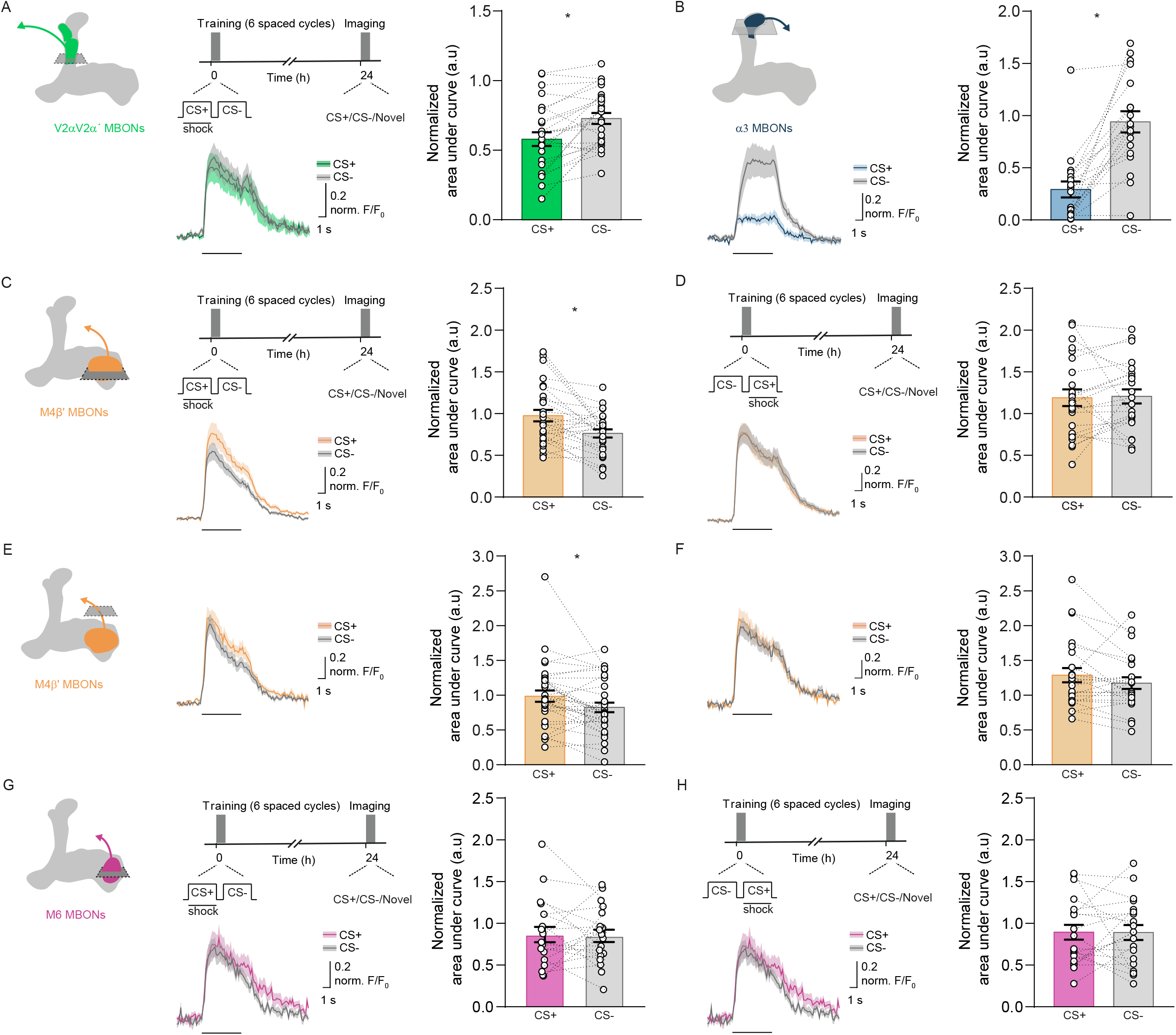
Parallel aversive and safety memories can be recorded as depression of odor-specific responses in corresponding MBONs. (**A** and **B**) Imaging planes in MBON-α2sc and MBON-α3 dendritic fields, training and imaging protocol. A reduced CS+ odor-evoked response was observed in both MBONs. (**C-F**) Imaging plane in MBON-β’2mp dendritic field (**C** and **D**) or presynaptic terminals (**E** and **F**), training and imaging protocol. Spaced training significantly reduces CS- responses in MBON-β’2mp dendritic (**C**) and axonal fields (**E**), but not when CS+ follows CS- in spaced training (**D** and **F**). (**G** and **H**). Imaging plane in MBON-γ5β’2a dendritic field, training and imaging protocol. Spaced training does not change the odor-evoked responses in MBON-γ5β’2a. CS+ data corresponds to average of data in which 50% of the trials used MCH as CS+ and 50% of were OCT CS+. Same applies for CS-. Asterisks denote significant difference between averaged CS+ and CS- responses. See also Table S1 and Figure S2.

We next tested for evidence of a CS- approach memory by recording odor-evoked responses in the MBON-β’2mp and MBON-γ5β’2a dendrites. Since our earlier experiments in **Figure 1** indicated that the order of CS+ and CS- presentation was of particular importance to generate CS- approach memory, we compared odor-evoked MBON responses from flies spaced trained with CS+/CS- ordered trials to those from flies spaced trained with CS-/CS+ trials. A reduced CS- odor-evoked response was measured in the dendritic and axonal fields of the β’2mp MBON (**Figure 3C and 3E**), in CS+/CS- space trained flies, but not when trials were reversed to CS-/CS+ (**Figure 3D and 3F**). No significant change in odor-evoked responses was measured in the MBON-γ5β’2a dendritic field 24 h after spaced training with either CS+/CS- (**Figure 3G and 3H**) or CS-/CS+ ordered trials (**Figure S2A and S2B**). A reduced response to the CS- in MBON-β’2mp dendrites after spaced training (only with CS+/CS- order trials) could, at least partially, contribute to the conditioned approach to the CS-.

### Aversive spaced training induces region-specific plasticity in the dendrites of γ3, γ3β’1 MBONs

The γ3 and γ3β’1 MBONs have also been implicated in aversive LTM (Wu et al., 2017). Since their dendrites occupy MB compartments that are innervated by PAM DANs, we used Ca^2+^ imaging to test for odor-evoked physiological correlates of CS- approach memory in the dendritic fields of these MBONs. The MB110C split-GAL4 driver expresses GCaMP in both γ3 and γ3β’1 MBONs. After spaced training, we observed strikingly different responses in the γ3 and β’1 dendritic fields, which are innervated by distinct PAM DANs. Recordings in the β’1 region revealed a reduced odor-evoked response to the CS- after spaced training (**Figure 4A**), but not when the CS+ and CS- order was reversed (**Figure 4B**). In contrast, a reduced odor-evoked response to the CS+ was observed in the γ3 dendrites, after spaced training but irrespective of the order of CS+ and CS- presentation (**Figure 4C and 4D**). Interestingly, recording in the presynaptic terminals of the γ3, γ3β’1 MBONs after training (**Figure 4E and 4F**) suggests that these neurons integrate the plasticity formed in the γ3 and β’1 dendritic regions. The relative decreased CS- response in the β’1 dendrite appears to be nullified when integrated with the relative decrease of the CS+ response in the γ3 dendritic portion. A trend for a reduced odor-evoked response to the CS+, which did not reach significance with this sample number, only remained when the flies were trained with reversed CS+ then CS- ordered trials (**Figure 4F**).

**Figure 4.**
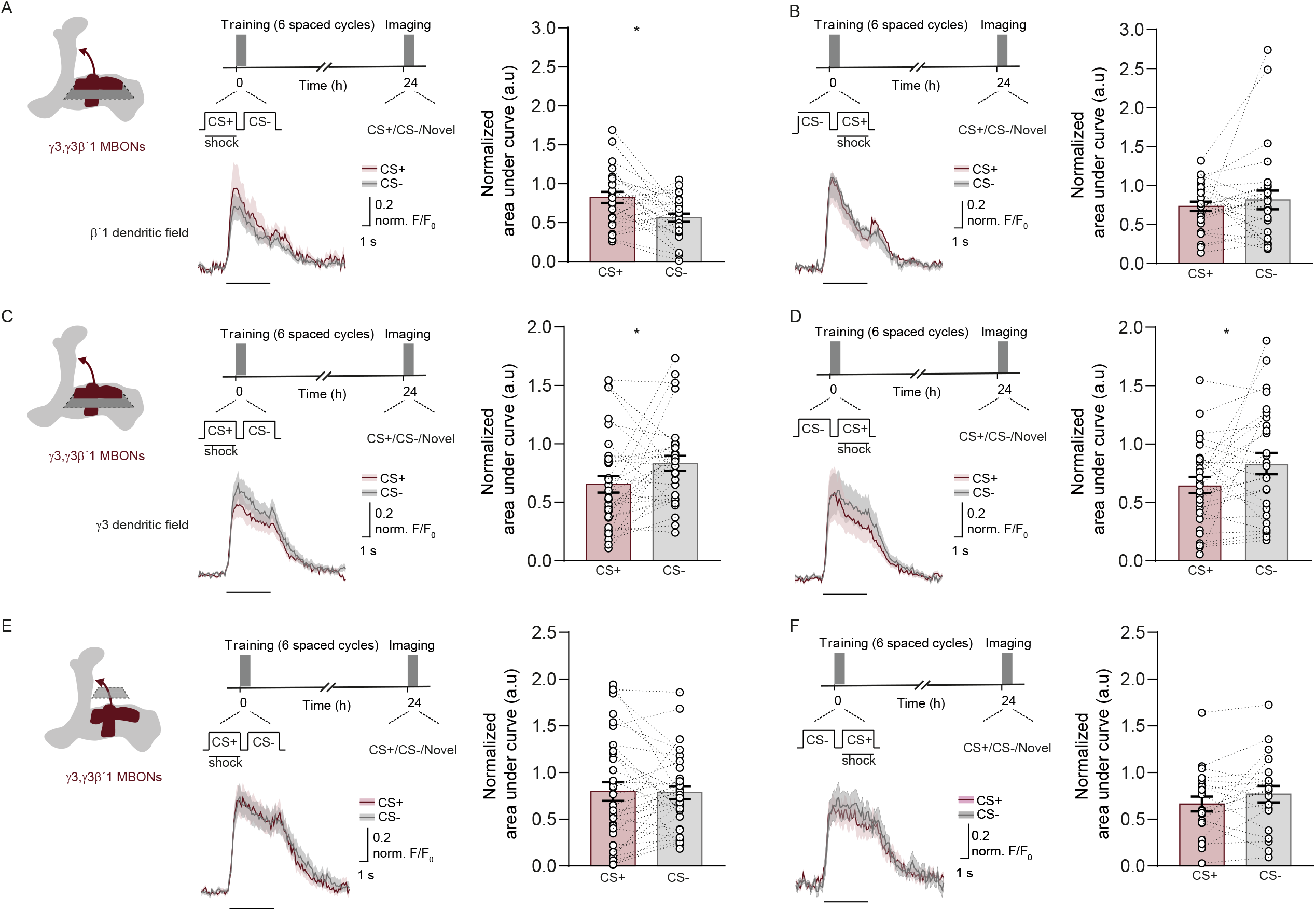
Spaced training induces region-specific plasticity of γ3, γ3β’1 MBON responses. (**A-F**) Measuring odor responses in γ3, γ3β’1 MBONs. (**A** and **B**) Imaging plane for β’1 region of γ3, γ3β’1 MBON dendritic field, training and imaging protocol. (**A**) Spaced training significantly reduces CS- responses in β’1, but not with the reverse protocol (**B**) where CS- precedes CS+ in each trial. (**C** and **D**). Imaging plane for γ3 region of γ3, γ3β’1 MBON dendritic field, training and imaging protocols. (**C**) Spaced training significantly reduces CS+ responses in γ3. (**D**) The reverse protocol also reduces CS+ responses in γ3. (**E** and **F**) Imaging plane in presynaptic terminals of γ3, γ3β’1 MBONs, training and imaging protocols. (**E**) Spaced training did not significantly alter odor-evoked responses in the γ3, γ3β’1 presynaptic terminals (**F**) A trend for a reduced response for the CS+ was observed, following the reverse protocol. CS+ data correspond to average of data in which 50% of trials used MCH as CS+, other 50% OCT was CS+. Same applies for CS- odor. Asterisks, significant difference between averaged CS+ and CS- responses. See also Table S1.

### Distributed plasticity is required for memory after spaced training

Finding depression of CS- odor-evoked responses in MBON-β’2mp (**Figure 3D**) and the β’1 tuft of the MBON-γ3, γ3β’1 dendritic field (**Figure 4A and 4C**), and decreased CS+ responses in α2sc- and α3-MBON dendrites (**Figure 3A and 3B**), suggests roles for the related DANs in LTM. We therefore used DAN specific control of UAS-*Shi*^*t*s1^ to test the importance of each site of plasticity for CS+ and CS- memories following spaced training.

The α2sc and α3 MBON compartments on the MB vertical lobe are innervated by aversively reinforcing DANs from the paired posterior lateral 1 (PPL1) cluster (**Figure 5A**). As expected, blocking output from PPL1-DANs using UAS-MB504B driven UAS-*Shi*^ts1^ severely impaired LTM performance when flies were tested for preference between CS+ and CS- odors (**Figure 5A**; one-way ANOVA: p<0.01, n=11). PPL1-DAN block specifically impaired CS+ aversive memory. No performance was observed when flies were tested CS+ *vs*. novel odor (**Figure 5B**; one-way ANOVA: p<0.05, n=7), whereas significant approach remained when flies were tested CS- *vs*. novel odor (**Figure 5C**; one-way ANOVA: p>0.05, n=8).

**Figure 5.**
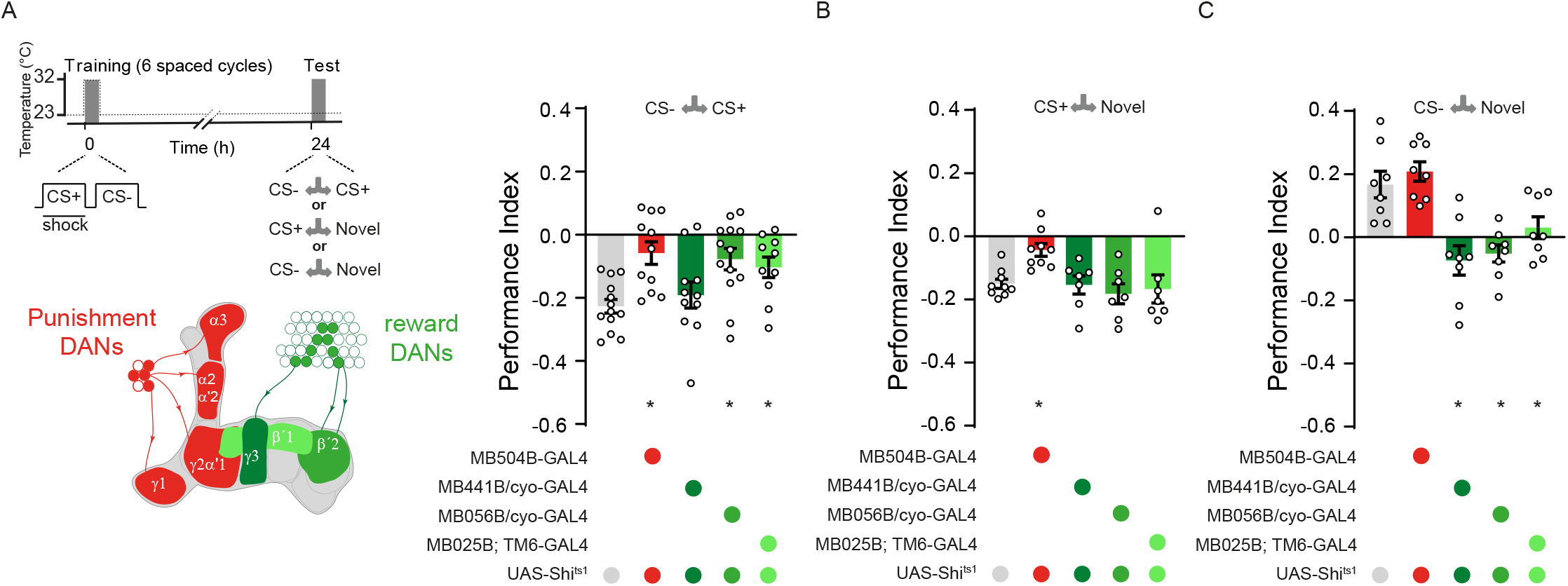
Blocking specific dopaminergic neurons during spaced training localises the discrete sites of aversive and safety memory. (**A**) Left: protocol with temperature shifting (dashed line) and color coded illustration of DANs labelled with each GAL4. Right: blocking PPL1-DANs during spaced training with MB504B-GAL4; UAS-*Shi*^ts1^ impaired 24 h memory. Performance was similarly impaired with PAM-β’2mp (MB056B-GAL4) or PAM-β’1 (MB025B-GAL4) block. Blocking PAM-γ3 (MB441B-GAL4) DANs did not impair 24 h performance. (**B**) Testing flies preference between CS+ and novel odor only revealed a significant impairment with PPL1-DAN block. (**C**) Individually Blocking PAM-β’2mp (MB056B-GAL4), PAM-β’1 (MB025B-GAL4) and PAM-γ3 (MB441B-GAL4) impaired preference CS- and novel odor, whereas blocking PPL1 DANs had no effect. Asterisks denote significant differences. Data are mean ± SEM. Individual data points displayed as dots.

We next tested the importance of DAN-directed plasticity in MBON-β’2mp and the two separable regions of the MBON-γ3,γ3β’1 dendritic fields for LTM formation. Blocking the PAM-β’2mp or PAM-β’1 DANs during training significantly impaired LTM performance when flies were tested for CS+ *vs*. CS- preference (**Figure 5A**; β’2mp, MB056B; one-way ANOVA: p<0.01, n=13. β’1, MB025B; one-way ANOVA: p<0.05, n=10). Defective performance could be specifically attributed to the CS- memory. Performance was unaffected if flies were tested CS+ *vs*. novel odor (**Figure 5B**; one-way ANOVA: p>0.05, n=7) whereas no significant performance was evident when flies were tested CS- vs. novel (**Figure 5C**; β’2; one-way ANOVA: p<0.001, n=8; β’1; one-way ANOVA: p<0.05, n=8). Blocking PAM-γ3 DANs revealed a surprisingly specific defect. LTM performance was not significantly impaired when flies were tested for CS+ vs. CS- (**Figure 5A**; MB441B; one-way ANOVA: p>0.05, n=11) or CS+ vs. novel (**Figure 5B**; one-way ANOVA: p>0.05, n=7). However, when PAM-γ3 blocked flies were tested for preference between CS- and novel odor no CS- approach was observed (**Figure 5C**; one-way ANOVA: p<0.001, n=8). Together these manipulations demonstrate roles for PPL1 DANs in plasticity representing the aversive CS+ memory and the PAM-β’2mp, PAM-β’1 and PAM-γ3 DANs for the CS- approach memory.

## Discussion

The memory performance gain from spacing learning sessions has intrigued scientists for over a century. Early work using *Drosophila* demonstrated that spaced training produced protein synthesis dependent “aversive LTM”, whereas massed training did not (Tully et al., 1994; Isabel et al., 2004). Many subsequent studies have compared memory performance after spaced training to that following massed training. We found that flies learn additional safety information for the CS- odor when subjected to spaced training. Parallel complementary CS+ aversive and CS- approach memories therefore account for the 24 h discriminative odor preference observed after differential spaced training. In contrast, flies only form an avoidance memory for the previously shock-paired odor following massed training. To our surprise, *radish* mutant flies did not form the CS+ aversive memory after spaced training yet their CS- memory appeared unaffected. In contrast, CXM feeding abolished the CS- memory, but the CS+ memory was not significantly reduced. Using previous operational definitions (Tully et al., 1994), these data suggest that the CS- memory is protein synthesis dependent LTM whereas the CS+ component is ARM. It is therefore important to rethink the many prior studies that have assumed to be measuring only avoidance of the CS+ following spaced training.

### Relief, or safety memory?

The valence of olfactory memories can be reversed from aversive to appetitive by changing the relative timing of odor and reinforcement presentation during training (Tanimoto et al., 2004; König et al., 2018). If shock, or artificial DAN activation is presented <45 seconds before odor presentation flies form an appetitive ‘relief memory’ for that odor (Tanimoto et al., 2004; Aso and Rubin, 2016; Handler et al., 2019). Experiments with artificial DAN activation suggest that relief learning is represented by dopamine potentiating an MBONs response to the conditioned odor (Handler et al., 2019; although see König et al., 2018). If spaced training utilizes the same relief from punishment mechanism as that in Handler et al. (2019), the CS- approach memory would be coded as potentiation of the same connections as those coding CS+ avoidance as a depression. However, we observed co-existence of aversive and approach memories at different places in the MBON network. Our data instead indicate that CS- approach is coded by specific appetitively reinforcing DANs that direct depression of KC outputs onto the corresponding MBON dendrites. We therefore propose that CS- approach after spaced training reflects the fly having formed a ‘safety memory’ for the CS-, rather than the CS- having been associated with the cessation of punishment. Massed training shortens the period of safety after each CS- presentation and does not form a CS- approach memory. In rodents, the neural underpinnings for relief learning and safety learning are also different (Mohammadi et al., 2014).

### Different MBONs guide CS+ and CS- performance after spaced training

Aversive LTM performance, after spaced training, is largely considered to rely on αβ KCs (Isabel et al., 2004), and to be retrieved via α2sc (MB-V2) MBONs (Séjourné et al., 2011; Bouzaiane et al., 2015). However, others have indicated that the network properties are more distributed and that output from γ3, 3β’1 and α3 MBONs is required to retrieve aversive LTM (Pai et al., 2013; Wu et al., 2017). Our work here suggests that there are likely to be different reasons for observing impairments when blocking these MBONs while testing 24 h memory following spaced training. Consistent with prior work (Séjourné et al., 2011) we identified depression of CS+ responses in α2sc-MBONs 24 h after spaced training. Depression of α2sc-MBONs neural responses therefore are critical for flies to express CS+ avoidance. We also uncovered very strong depression of CS+ responses in MBON-α3 after spaced training. The role for α3 MBONs has been disputed (Pai et al., 2013; Plaçais et al., 2013). At this point we cannot reconcile the differences between the three studies, other than perhaps the number of training trials, strength of reinforcement, and the relative hunger state of the flies, some of which was also proposed by Plaçais et al. (2013). We also note that many recent studies train flies using robotic control where flies remain in the same training tube for the entire session. In contrast, earlier studies and our experiments reported here, utilised manual training where flies are transferred from the training chamber context between trials. Nevertheless, our data here suggest that α2sc and α3 MBONs house plasticity relevant for the expression of CS+ aversive memory.

Bouzainne et al., (2015) reported that MBON-β’2mp and MBON-γ5β’2a (M4/6) are not required for retrieval of LTM, after spaced training. However, we found that appropriately ordered CS+/CS- spaced training trials induce a depression of CS- responses in dendrites of β’2mp-MBONs. In addition, we found that the PAM-β’2mp DANs are required to form the CS- approach memory. Our results therefore indicate a specific role for the β’2mp subcompartment of the β’2 MB zone and that MBON-β’2mp plasticity is required to express CS- approach memory, in agreement with findings from Wu et al. (2017).

The negative sign of odor response plasticity of α2sc, α3 and β’2mp MBONs makes intuitive sense with the known valence of these pathways. CS+ responses in approach-directing α2sc and α3 MBONs were depressed, which would favor odor avoidance. In contrast, depressing CS- responses to avoidance-directing β’2mp MBONs should promote odor approach.

### γ3β’1 MBONs compute and provide a measure of relative safety?

We also uncovered roles for PAM-γ3 and PAM-β’1 DANs and discovered traces of both the CS+ and CS- memory when recording in the corresponding γ3 and γ3β’1 MBONs. MBON dendrites in the γ3 compartment showed a decreased response to the CS+, irrespective of the order of the CS+ and CS- training trials, consistent with the rules of forming an aversive CS+ memory. In contrast, responses to the CS- were decreased in the β’1 tuft of the γ3β’1 MBON dendrite, but only if flies were trained with CS+ then CS- ordered trials. Plasticity in β’1 of the γ3β’1 MBON therefore followed the order rule observed for conditioning CS- approach behaviour. Interestingly, recording in the axons of γ3 and γ3β’1 MBONs suggested that the CS+ and CS- plasticity cancel each other out. Regrettably, the split GAL4 used to drive GCaMP expression in γ3β’1 MBONs also labels γ3 MBONs. Therefore, although only the γ3β’1 MBONs have a dendrite in both the γ3 and β’1 compartments, we cannot at this stage be certain that the γ3β’1 MBONs alone integrate the CS+ and CS- memory traces.

To decipher the relative role of γ3 and β’1 plasticity we individually blocked output from PAM-γ3 and PAM-β’1 DANs during training and tested the resulting memories. The behavioral observations following PAM-γ3 block were particularly revealing. PAM-γ3 DANs respond to shock (Cohn et al., 2015) and their forced activation reinforces aversive memories (Yamagata et al., 2016). However, blocking PAM-γ3 DANs during spaced training did not impair CS+ avoidance and instead impaired CS- approach when flies were tested for CS- vs. novel odor preference. A CS- memory defect was also observed when blocking the appetitively reinforcing PAM β’1 DANs during training, although this manipulation also impaired CS+ vs. CS- performance. We therefore propose that γ3β’1 MBONs integrate the γ3 CS+ danger and β’1 CS- safety plasticity to compute a relative safety signal. The importance of which is only obvious when we remove aversive CS+ plasticity in γ3, and thereby reveal the behavioral consequence of unopposed CS- plasticity in the β’1 region of the γ3β’1 MBON. Since MBON-γ3 and MBON-γ3β’1 are GABAergic (Aso et al., 2014a), CS+ then CS- plasticity evoked by spaced training sequentially alters the level of inhibition imposed on their unknown downstream target neurons.

### A subset of rewarding DANs code learned safety

Our results here demonstrate that rewarding DANs reinforce the delayed recognition of safety. Formation of the CS- approach memory requires the appetitively reinforcing PAM-β’2mp and PAM-β’1 DANs and surprisingly, the aversively reinforcing PAM-γ3 DANs. As noted above PAM-γ3 DANs are shock activated (Cohn et al., 2015) and likely provide an aversive teaching signal (Yamagata et al., 2016) that directs CS+ plasticity in the γ3 region of the MBON-γ3β’1 dendrites. However, we are currently unsure what triggers the appetitively reinforcing PAM-β’2mp and PAM-β’1 DANs so that they code an approach memory for the CS- as decreased CS- activation of MBON-β’2mp and the β’1 region of MBON-γ3β’1, respectively. Blocking output from the PAM-β’2mp, PAM-β’1 or PAM-γ3 DAN, that are presumably responsible for each part of the LTM correlated plasticity, reveals that they are required during training to form CS- approach memories. Blocking most PAM-DANs further restricted their essential function to during CS- presentation in each spaced training trial, suggesting that they could be driven by the CS- odor. However, CS- memory formation requires that each CS+ exposure must precede each CS- in each training trial. Therefore PAM-DANs also have to somehow register a temporally locked negatively reinforced CS+ reference to be able to classify the following CS- as safe. Lastly, repetition is a necessary element of triggering DANs to code safety.

It is conceivable that formation of long-term CS+ and CS- memories is orchestrated by aversive reinforcement signals provided by the PPL1-γ1pedc, (MP1) and PPL1-γ2α’1 (MV1) DANs in each shock-paired CS+ trial (Claridge-Chang et al., 2009; Aso et al., 2010, 2014a; Plaçais et al., 2012; Aso and Rubin, 2016). The PPL1-γ1pedc DAN codes learning by depressing odor specific input to the feedforward GABAergic MBON-γ1pedc>αβ (MVP2) (Hige et al., 2015; Perisse et al., 2016). Although MVP2 output is only required for the expression of short-term aversive memory, the plasticity remains for several hours (Perisse et al., 2016). Each shock-reinforced odor trial therefore changes the state of the rest of the MBON/DAN network for the subsequent exposures and reinforced trials. This has been proposed to release the PPL1-α’2α2 DANs so that they can reinforce LTM at the KC-MBON-α2sc junction (Awata et al., 2019). A similar release from inhibition of the PPL1-α3, PAM-β’2mp, PAM-β’1 and PAM-γ3 DANs could account for our spaced training-dependent plasticity at MBON-α3 and prime the PAM-β’2mp and PAM-β’1 to reinforce the CS- memory.

Reinforcing PAM DANs have also been implicated in memory formation with sugar (Burke et al., 2012; Liu et al., 2012), water (Lin et al., 2014; Shyu et al., 2017) and alcohol reward (Scaplen et al., 2019), relative shock (Perisse et al., 2013), absence of expected shock (Felsenberg et al., 2018), and following courtship (Keleman et al., 2012). In addition, they provide the control of state-dependent memory expression (Senapati et al., 2019) and unlearned behavioral responses to volatile cues (Lin et al., 2014; Lewis et al., 2015). In some cases these processes clearly involve different DANs whereas in others they appear to involve DANs that innervate the same MB compartments. More refined tools, connectomics (Zheng et al., 2018) and experiments should help reveal the full extent of functional heterogeneity.

### Ubiquity and utility of parallel memories

Fly behavioral performance has previously been shown to be determined by combining either supporting or conflicting memories. When differentially conditioned by pairing one odor with shock and the other with sugar flies show additive initial performance to that observed if only one of the two odors had been reinforced (Tempel et al., 1983). This situation resembles that described here after spaced training but the second odor is explicitly unpaired, rather than sugar reinforced, and the additive performance emerges from complementary LTM. With the benefit of retrospect, and as nicely discussed before (Schleyer et al., 2018), it makes intuitive sense that over repetitive spaced trials flies learn ‘where the punishment is, and where it is not’. These parallel memories make it easier for flies to distinguish between the two odors when tested together (Barth et al., 2014).

In contrast, flies simultaneously form parallel competing memories when trained with bitter tainted sugar and their performance switches from aversion to approach over time, as dictated by the superior persistence of the nutrient-dependent sugar memory (Das et al., 2014). A similar time-dependent behavioral transition is evident when flies are trained with alcohol reinforcement (Kaun et al., 2011). A competition between memories of opposing valence also underlies the extinction of both appetitive and aversive memories (Felsenberg et al., 2017, 2018). However, opposing extinction memories are sequentially formed, being reinforced by the absence of an expected outcome, rather than explicit pairing. In these cases forming parallel memories reduces the certainty of odor choice.

Together these studies suggest that forming parallel memories in different places is a general feature of the *Drosophila* MBON network that summates experience over time to optimize the expression of learned behavior.

## Acknowledgments

We thank G. Rubin, FlyLight, B. Dickson and the Bloomington Stock Center for flies. We are grateful to Johannes Felsenberg, Vincent Croset, Emmanuel Perisse, Geraldine Wright and other members of the Waddell group for discussion and comments on the manuscript. S.W. is funded by a Wellcome Principal Research Fellowship (200846/Z/16/Z), Gatsby Charitable Foundation (GAT3237), ERC Advanced Grant (789274) and Bettencourt–Schueller Foundations.

## Author contributions

Conceptualization P. F. J., S.W., Methodology P.F.J., S.W., Investigation P.F.J. Resources S.W. Writing S.W., P.F.J. Supervision S.W. Funding Acquisition S.W.

## Star Methods

### Key Resource Table

| REAGENT or RESOURCE | SOURCE | IDENTIFIER |
| --- | --- | --- |
| **Experimental Models: Organisms/Strains** |  |  |
| *Drosophila*: MB110C-Gal4 | Bloomington Drosophila Stock Center;<br>(Aso et al., 2014a, 2014b) | RRID:BDSC_68262 |
| *Drosophila*: R66C08-Gal4 | Bloomington Drosophila Stock Center;<br>(Owald et al., 2015) | RRID:BDSC_49412 |
| *Drosophila*: R39A05-Gal4 | Bloomington Drosophila Stock Center;<br>(Jenett et al., 2012) | RRID:BDSC_50033 |
| *Drosophila*: R71D08-Gal4 | Bloomington Drosophila Stock Center;<br>(Jenett et al., 2012) | RRID:BDSC_61645 |
| *Drosophila*: G0239-Gal4 | Bloomington Drosophila Stock Center;<br>(Pai et al., 2013) | RRID:BDSC_12639 |
| *Drosophila*: R58E02-Gal4 | Bloomington Drosophila Stock Center;<br>(Liu et al., 2012) | RRID:BDSC_41347 |
| *Drosophila*: MB504B-Gal4 | Bloomington Drosophila Stock Center;<br>(Aso et al., 2014a, 2014b) | RRID:BDSC_68329 |
| *Drosophila*: MB056B-Gal4 | Bloomington Drosophila Stock Center;<br>(Aso et al., 2014a, 2014b) | RRID:BDSC_68276 |
| *Drosophila*: MB441B-Gal4 | Bloomington Drosophila Stock Center;<br>(Aso et al., 2014a, 2014b) | RRID:BDSC_68251 |
| *Drosophila*: MB025B-Gal4 | Bloomington Drosophila Stock Center;<br>(Aso et al., 2014a, 2014b) | RRID:BDSC_68299 |
| *Drosophila*: *radish* mutant | (Folkers et al., 1993) | N/A |
| *Drosophila*: UAS-GCaMP6m | Bloomington Drosophila Stock Center;<br>(Chen et al., 2013) | RRID:BDSC_42748 |
| *Drosophila*: UAS-GtACR1 | (Mohamed et al., 2017) | N/A |
| *Drosophila*: UAS-*Shi*<sup>ts1</sup> | (Kitamoto, 2001) | N/A |
| **Chemicals, Peptides, and Recombinant Proteins** |  |  |
| N-Tris | Sigma-Aldrich | Cat#T5691 |
| NaCl | Sigma-Aldrich | Cat#S7653 |
| KCl | Sigma-Aldrich | Cat#P9333 |
| NaHCO<sub>3</sub> | Sigma-Aldrich | Cat#S6297 |
| NaH<sub>2</sub>PO<sub>4</sub> | Sigma-Aldrich | Cat#S8282 |
| CaCl<sub>2</sub> | Sigma-Aldrich | Cat#21115 |
| MgCl<sub>2</sub> | Sigma-Aldrich | Cat#M1028 |
| Trehalose | Sigma-Aldrich | Cat#T9531 |
| Glucose | Sigma-Aldrich | Cat#G7528 |
| Sucrose | Sigma-Aldrich | Cat# S0389 |
| Mineral Oil | Sigma-Aldrich | Cat#M5904 |
| 4-methylcyclohexanol (98%) | Sigma-Aldrich | Cat#218405 |
| 3-octanol (99%) | Sigma-Aldrich | Cat#153095 |
| Isopentyl acetate (99%) | Sigma-Aldrich | Cat#306967 |
| Cycloheximide | Sigma-Aldrich | Cat#7698 |
| **Software and Algorithms** |  |  |
| Fiji | NIH; (Schindelin et al., 2012) | http://fiji.sc/ |
| Matlab R2017b | The Mathworks, Natick, MA | https://www.mathworks.com/products/matlab.html |
| GraphPad Prism 7 | GraphPad Software, La Jolla, CA | https://www.graphpad.com/scientific-software/prism/ |
| Adobe Illustrator CC | Adobe Systems, San Jose, CA | https://www.adobe.com/uk/products/illustrator.html |
| ScanImage 3.8 software | (Pologruto et al., 2003) | https://vidriotechnologies.com/ |

### CONTACT FOR REAGENT AND RESOURCE SHARING

Further information and requests for resources and reagents should be directed to and will be fulfilled by Scott Waddell

### EXPERIMENTAL MODEL AND SUBJECT DETAILS

#### Fly strains

All *Drosophila melanogaster* strains were reared at 25°C and 40-50% humidity on standard cornmeal-agar food in 12:12 h light:dark cycle. Flies from the wild-type (WT) Canton-S and mutant *radish* (*rad*) (Folkers et al., 1993) strains were used. Transgenes were expressed with previously described GAL4 lines: R58E02-GAL4 (Liu et al., 2012), MB110C-GAL4, MB504B-GAL4, MB056B-GAL4, MB056B-GAL4, MB441B-GAL4 and MB025B-GAL4 (Aso et al., 2014a, 2014b), R66C08-GAL4 (Owald et al., 2015), R39A05-GAL4 and R71D08 (Jenett et al., 2012); G0239-GAL4 (Pai et al., 2013). For behavioral experiments UAS-*Shi*^ts1^ (Kitamoto, 2001) and GtACR1 (Mohamed et al., 2017) were expressed under the control of the respective GAL4–line. For the imaging experiments UAS-GCaMP6m (Chen et al., 2013) was expressed with the respective GAL4.

### METHOD DETAILS

#### Behavioral experiments

Male flies from the GAL4 lines were crossed to UAS-*Shi*^ts^ or GtACR1 females and 4 to 9-day-old mixed-sex progeny were tested together in all experiments. For WT, CXM fed and *rad* experiments flies with 4 to 9-day-old mixed-sex progeny were used. Approximately 80 - 100 flies were placed in a 25 ml vial containing standard food and a 20 × 60 mm piece of filter paper for 14–22 hours before behavioral experiments, except where noted. Odors used in all experiments were 4-methylcyclohexanol (MCH), 3-octanol (OCT) and isoamyl acetate (IAA) diluted in mineral oil. An odor dilution of ~1:10^3^ for all experiments (specifically, 8-12 μl OCT, 8-9 μl MCH or 16-18 μl IAA in 8 ml mineral oil). All experiments were performed at 23°C, except where noted, and 55-65% relative humidity.

For experiments involving neuronal blockade with *Shi^ts1^*, the time courses of the temperature shifts are provided alongside each graph of memory performance. Flies were transferred to the restrictive temperature (32°C) 30min before the targeted time, except where noted, to allow for acclimatization to the new temperature. Prior to optogenetic experiments all flies were housed on standard cornmeal food with 1mM retinal for 3 days.

Aversive olfactory conditioning in the T-maze was conducted as previously described (Tully and Quinn, 1985; Perisse et al., 2016). Groups of flies were trained with either one cycle of aversive training, six consecutive cycles (massed training) or six cycles spaced by 15 minute inter-trial intervals (spaced training). After each cycle when spaced training flies were transferred from the training tube back into their starter vial, until the start of the next cycle. Except where noted, during each cycle of training, flies were exposed to a first odor for 1 min (the conditioned stimulus+, CS+) paired with twelve 90 V electric shocks at 5 s intervals. Following 45 s of clean air, a second odor (the conditioned stimulus-, CS-) was presented for 1 min without shock. Flies were kept in food vials at 23°C between training and test. Memory was subsequently assessed 24 h after training by testing flies for their odor-preference between the CS- and the CS+ or the CS+ or CS- versus novel odor in a T-maze (2 min, in darkness). The testing odors were always MCH and OCT, where a novel odor was applied during the test IAA was used either as CS+ or CS-. Performance index was calculated as the number of flies in the CS+ arm minus the number in the CS- arm, divided by the total number of flies (Tully and Quinn, 1985). When the performance was tested against a novel odor the performance index was calculated as the number of flies in the CS+ or CS- arm minus the number in the Novel arm, divided by the total number of flies. MCH and/or OCT and/or IAA, were alternately used as CS+ or CS- and a single sample, or n, represents the average performance score from two reciprocally trained groups.

#### CXM feeding

WT flies were fed with cycloheximide (CXM) for 12-16 h prior to training as reported before (Tully et al., 1994; Yin et al., 1994). In brief, filter paper strips were soaked with 250 μl 5% glucose solution that was laced with 35 mM CXM. For controls flies, the filter paper strips were soaked with 250 μl 5% glucose solution. Flies were then transferred to the training apparatus and subject to spaced training. They were then transferred to test tubes containing filter paper strips soaked with 5% glucose during the 24 h retention interval before testing.

#### Two-photon Calcium Imaging

3-8 day old flies were imaged 23-25 h after aversive conditioning. Flies were trained as described above. Imaging experiments were performed essentially as described previously (Owald et al., 2015; Perisse et al., 2016; Felsenberg et al., 2018). In brief, flies were immobilized on ice and mounted in a custom made chamber allowing free movement of the antennae and legs. The head capsule was opened under room temperature carbogenated (95% O_2_, 5% CO_2_) buffer solution (103 mM NaCl, 3 mM KCl, 5mM N-Tris, 10 mM trehalose, 10 mM glucose, 7mM sucrose, 26 mM NaHCO_3_, 1mM NaH_2_PO_4_, 1.5 mM CaCl_2_, 4mM MgCl_2_, osmolarity 275 mOsm, pH 7.3) and the fly, in the recording chamber, was placed under the Two-Photon microscope (Scientifica). A constant air stream, carrying vapor from mineral oil solvent (air) was applied. GCaMP responses to the CS+, the CS- and a novel odor were measured in the relevant MBONs. Flies were sequentially exposed to the CS+, CS- and a novel odor, isoamyl acetate (IAA; 1:10^3^ odor concentration) for 5 s. Each odor presentation was followed by 30 s of air. To image the dendritic field and axonal segments of MBON-γ3,γ3β’1, the axonal segments of the MBON-β’2mp and MBON-γ5β’2a processes, the dendritic field of MBON-α2sc and MBON-α3 one hemisphere of the brain was randomly selected. To measure responses in the MBON-β’2mp and MBON-γ5β’2a dendrites, signals were simultaneously acquired from both hemispheres and averaged responses were analyzed.

Fluorescence was excited using ~140 fs pulses, 80 MHz repetition rate, centered on 910 nm generated by a Ti-Sapphire laser (Chameleon Ultra II, Coherent). Images were acquired with a Two-Photon microscope with a 403, 0.8 NA water-immersion 40X objective, controlled by ScanImage 3.8 software(Pologruto et al., 2003). Odors were delivered using a custom-designed system (Shang et al., 2007).

For analysis, two-photon fluorescence images were manually segmented using Fiji (Schindelin et al., 2012). Movement of the animals was small enough such that images did not require registration. For subsequent quantitative analyses, custom Fiji and MATLAB scripts were used. The baseline fluorescence, F_0_, was defined for each stimulus response as the mean fluorescence F from 2 s before and up to the point of odor presentation. F/F_0_ accordingly describes the fluorescence relative to this baseline. The area under the curve (AUC) was measured as the integral of F/F_0_ during the 5 s odor stimulation. To account for variance between individual flies, the responses of the CS+ and CS- were normalized to the response to IAA. Each AUC was divided by the IAA AUC from the respective trial and individual fly.

### QUANTIFICATION AND STATISTICAL ANALYSIS

Statistical analyses were performed in GraphPad Prism. All behavioral data were analyzed with an unpaired t-test or an one-way ANOVA followed by Tukey’s multiple comparisons test as post hoc test. No statistical methods were used to predetermine sample size. For the imaging experiments normalized responses were compared by a paired t-test for normally distributed data, otherwise a Wilcoxon matched-pairs signed rank test was used for non-Gaussian distributed data. The respective statistical tests used, the n numbers and the p values for the imaging data can be found in Table S1.

### DATA AND SOFTWARE AVAILABILITY

Customized MATLAB and Fiji scripts used in this paper are available.

## Supplementary Figure Legends

**Figure S1.**
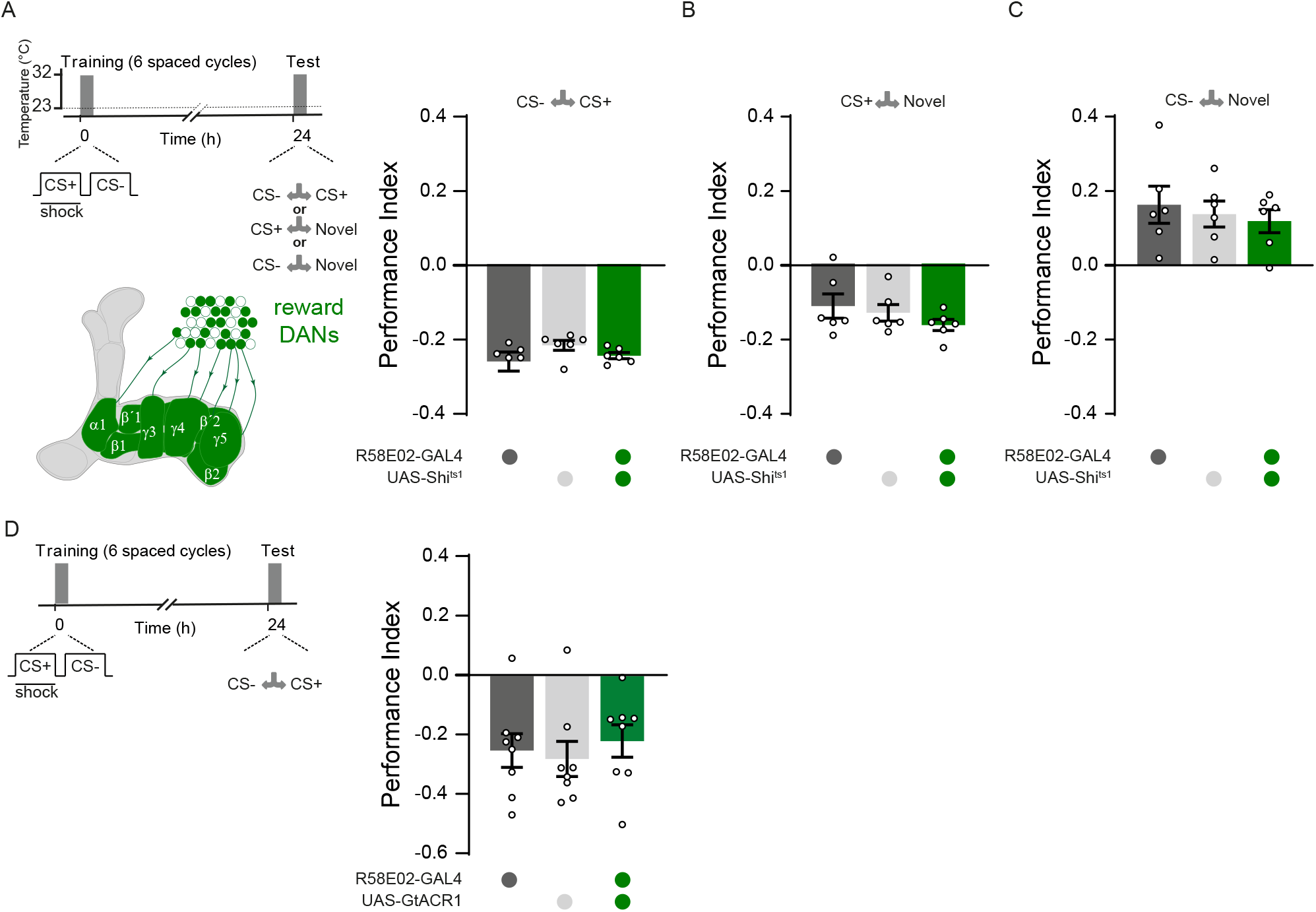
Temperature and light controls related to Figure 2. (**A**) Left: protocol and PAM/R58E02-GAL4 DAN schematic. Right: Expressing UAS-*Shi*^ts1^ in PAM DANs (R58E02-GAL4) does not disrupt 24 h memory performance after spaced training at permissive 23°C (one-way ANOVA: p>0.05; n=6). (**B**) Aversive memory for the CS+ is unaffected (one-way ANOVA: p>0.05; n=6). (**C**) Approach memory to CS- is unaffected (one-way ANOVA: p>0.05; n=6). (**D**) Left: protocol without presentation of green light. Right: UAS-GtACR1 expression in PAM DANs (R58E02-GAL4) does not disrupt 24 h memory performance after spaced training (one-way ANOVA: p>0.05; n=8)

**Figure S2.**
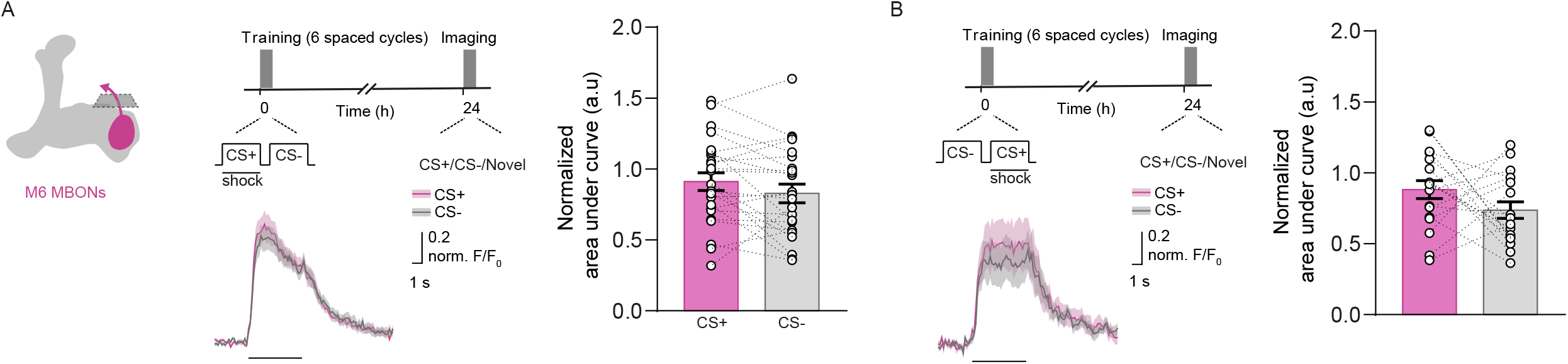
Odor responses in MBON-γ5β’2a are unchanged following spaced training. Imaging plane in presynaptic terminals of MBON-γ5β’2a, training and imaging protocols. (**A**) Spaced training and (**B**) the reversed protocol do not change odor-evoked responses in MBON-γ5β’2a. See also Table S1.

**Table S1.** Statistical details related to Figure 3, Figure 4 and Figure S2.

| Figure | Experiment | n | Normally distributed | Statistical test | p value |
| --- | --- | --- | --- | --- | --- |
| Figure 3A | R71D08-GAL4 after spaced training | 28 | yes | paired t-test | $p = 0.0002$ |
| Figure 3B | G0239-GAL4 after spaced training | 20 | no | Wilcoxon matched-pairs signed rank | $p < 0.0001$ |
| Figure 3C | R39A05-GAL4 after spaced training dendrites | 28 | yes | paired t-test | $p = 0.023$ |
| Figure 3D | R39A05-GAL4 after spaced training reverse order dendrites | 24 | yes | paired t-test | $p = 0.851$ |
| Figure 3E | R39A05-GAL4 after spaced training presynaptic terminals | 34 | yes | paired t-test | $p = 0.013$ |
| Figure 3F | R39A05-GAL4 after spaced training reverse order presynaptic terminals | 24 | yes | paired t-test | $p = 0.118$ |
| Figure 3G | R66C08-GAL4 after spaced training dendrites | 20 | yes | paired t-test | $p = 0.8591$ |
| Figure 3H | R66C08-GAL4 after spaced training reverse order dendrites | 20 | yes | paired t-test | $p = 0.9748$ |
| Figure 4A | MB110C-GAL4 after spaced training $\beta$'1 dendrites | 34 | yes | paired t-test | $p = 0.0006$ |
| Figure 4B | MB110C-GAL4 after spaced training reverse order $\beta$'1 dendrites | 30 | no | Wilcoxon matched-pairs signed rank | $p = 0.749$ |
| Figure 4C | MB110C-GAL4 after spaced training γ3 dendrites | 36 | no | Wilcoxon matched-pairs signed rank | $p = 0.011$ |
| Figure 4D | MB110C-GAL4 after spaced training reverse order γ3 dendrites | 30 | yes | paired t-test | $p = 0.04$ |
| Figure 4E | MB110C-GAL4 after spaced training presynaptic terminals | 34 | yes | paired t-test | $p = 0.899$ |
| Figure 4F | MB110C-GAL4 after spaced training reverse order presynaptic terminals | 22 | yes | paired t-test | $p = 0.14$ |
| Figure S2A | R66C08-GAL4 after spaced training presynaptic terminals | 24 | yes | paired t-test | $p = 0.0825$ |
| Figure S2B | R66C08-GAL4 after spaced training reverse order presynaptic terminals | 18 | yes | paired t-test | $p = 0.1615$ |

